## Supplementary figures for "Astrocytic modulation of population encoding in mouse visual cortex via GABA transporter 3 revealed by multiplexed CRISPR/Cas9 gene editing"

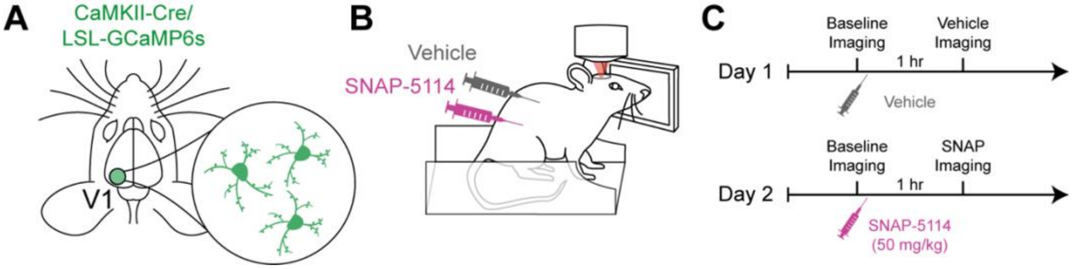


**Figure S1. Experimental design and timeline** (A) Transgenic mice expressing Cre-dependent GCaMP6s in excitatory neurons (CaMKII-Cre) had cranial windows implanted over V1 and were imaged 1-2 weeks later. (B) Neuronal responses to visual stimuli were imaged using two-photon microscopy in head-fixed mice. Following baseline imaging, either vehicle (5% DMSO in corn oil) or SNAP-5114 (50mg/kg) was injected i.p. followed by imaging the same field-of-view (FOV). (C) Timeline for imaging. Imaging after vehicle and SNAP-5114 treatment was performed on the same FOV on two different days. Both were compared to baseline imaging.

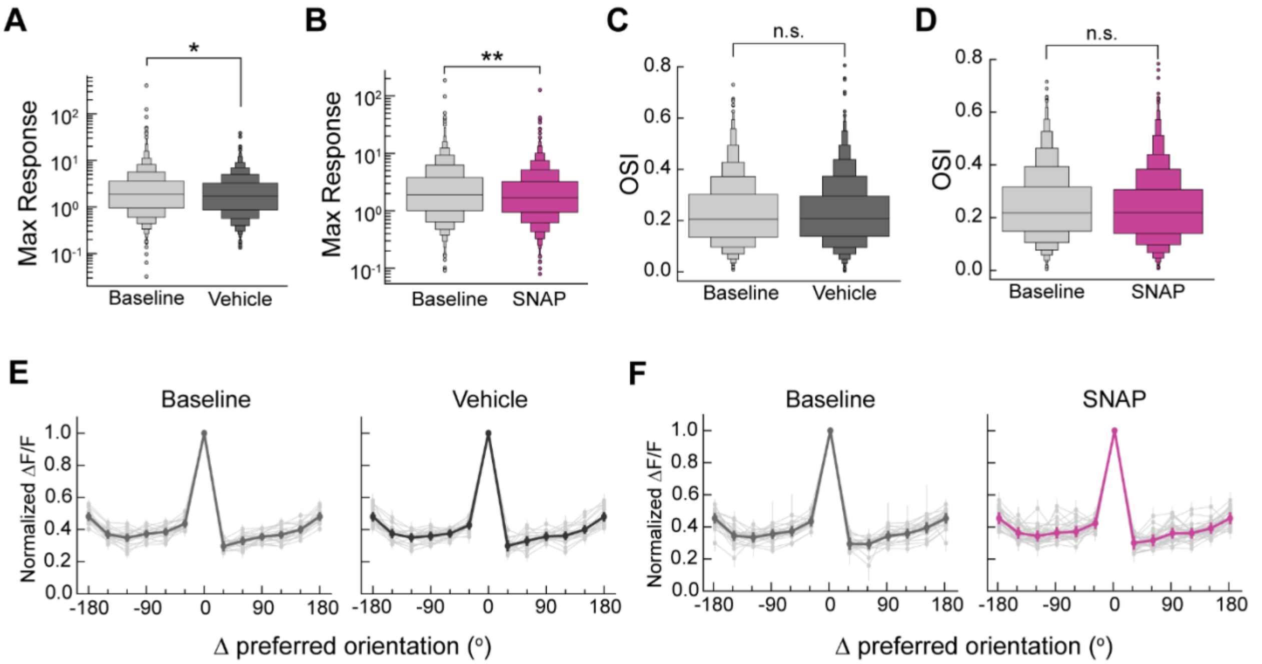


**Figure S2. Systemic administration of SNAP-5114 affects select V1 neuronal response properties** (A) Average maximum response magnitude of visually responsive neurons before and after vehicle administration. ($n_{baseline}$ = 1798 neurons, $n_{vehicle}$ = 1648 neurons, $n_{baseline}$ = 20 sessions, $n_{vehicle}$ = 20 sessions, *, $p < 0.05$, LME t-stats). (B) Same as A but before and after SNAP-5114 administration ($n_{baseline}$ = 1709 neurons, $n_{SNAP-5114}$ = 2002 neurons, $n_{baseline}$ = 25 sessions, $n_{SNAP-5114}$ = 38 sessions, **, $p < 0.01$, LME t-stats). (C) Orientation selectivity index (OSI) distribution of all visually responsive neurons before and after vehicle administration ($n_{baseline}$ = 1798 neurons, $n_{vehicle}$ = 1648 neurons, $n_{baseline}$ = 20 sessions, $n_{vehicle}$ = 20 sessions, n.s., $p = 0.359$, LME t-stats). (D) Same as C but before and after SNAP-5114 treatment ($n_{baseline}$ = 1709 neurons, $n_{SNAP-5114}$ = 2002 neurons, $n_{baseline}$ = 25 sessions, $n_{SNAP-5114}$ = 38 sessions, n.s., $p = 0.622$, LME t-stats). (E) Average tuning curves of visually responsive neurons per session (in lighter shade) before and after vehicle administration. Average tuning curve of all sessions is in bold ($n_{baseline}$ = 1798 neurons, $n_{vehicle}$ = 1648 neurons, $n_{baseline}$ = 20 sessions, $n_{vehicle}$ = 20 sessions, error bars = SEM). (F) Same as E but before and after SNAP-5114 treatment ($n_{baseline}$ = 1709 neurons, $n_{SNAP-5114}$ = 2002 neurons, $n_{baseline}$ = 25 sessions, $n_{SNAP-5114}$ = 38 sessions, error bars = SEM). All sessions were from 5 mice studied under vehicle and SNAP-5114 treatment conditions (see Figure S1).

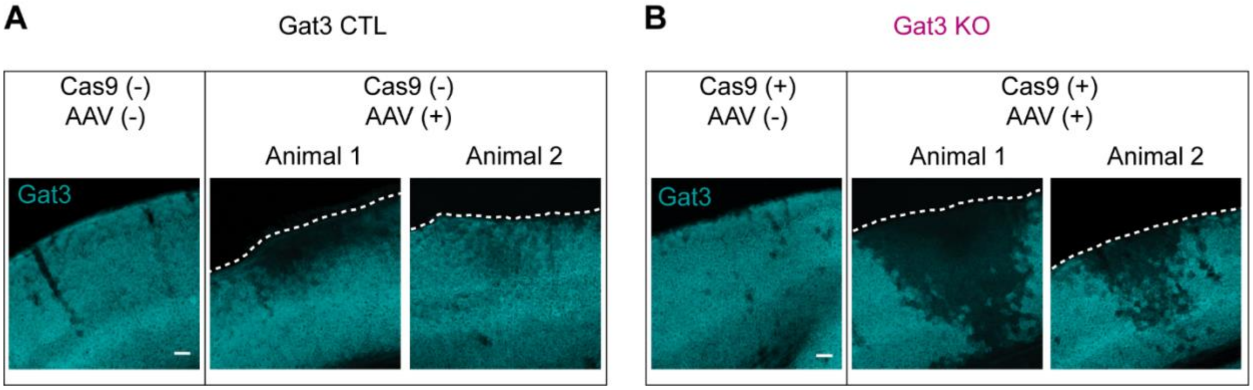


**Figure S3. Gat3 expression changes in control and KO brain slices** (A) Coronal slices of control animal brains. Left column: hemisphere with no virus injection. Right columns: hemispheres with virus injection from two different animals. Reduced superficial Gat3 expression is observed in control animals with variability between animals. Animal 1 represents the maximal effect of AAV injection on Gat3 expression and animal 2 represents a more typical effect (scale bar = 100 μm, applies to all images). (B) Same as A but for Gat3 KO animal brains. Compared to the injected hemispheres of control animals, the injected hemispheres of Gat3 KO animals showed a more severe reduction of Gat3 expression throughout the cortical layers (scale bar = 100 μm, applies to all images). White dotted lines indicate pial surface of the cortex.

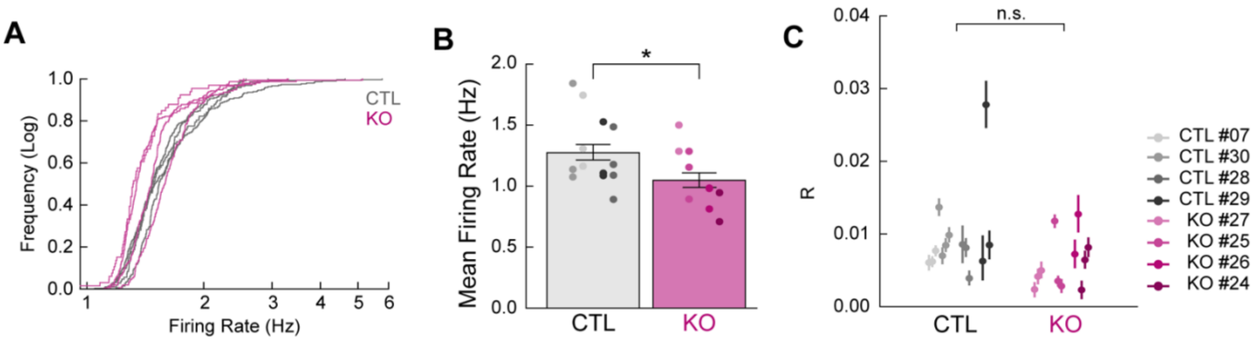


**Figure S4. Comparisons of average firing rates of single neurons and neuron-to-neuron pairwise correlation during spontaneous activity** (A) Empirical cumulative distribution functions plot of firing rates for each animal shown as individual traces ($n_{control}$ = 4 mice, $n_{Gat3\ KO}$ = 4 mice). (B) Average firing rates of neurons; different sessions per animal indicated by different colors ($n_{control}$ = 13 sessions, 4 mice, $n_{Gat3\ KO}$ = 11 sessions, 4 mice, *, $p < 0.05$, LME t-stats, error bars = SEM). (C) Average neuron-to-neuron pairwise correlation coefficient of neurons per session by animal ($n_{control}$ = 13 sessions, 4 mice, $n_{Gat3\ KO}$ = 11 sessions, 4 mice, n.s., $p = 0.224$, LME t-stats, error bars = SEM).

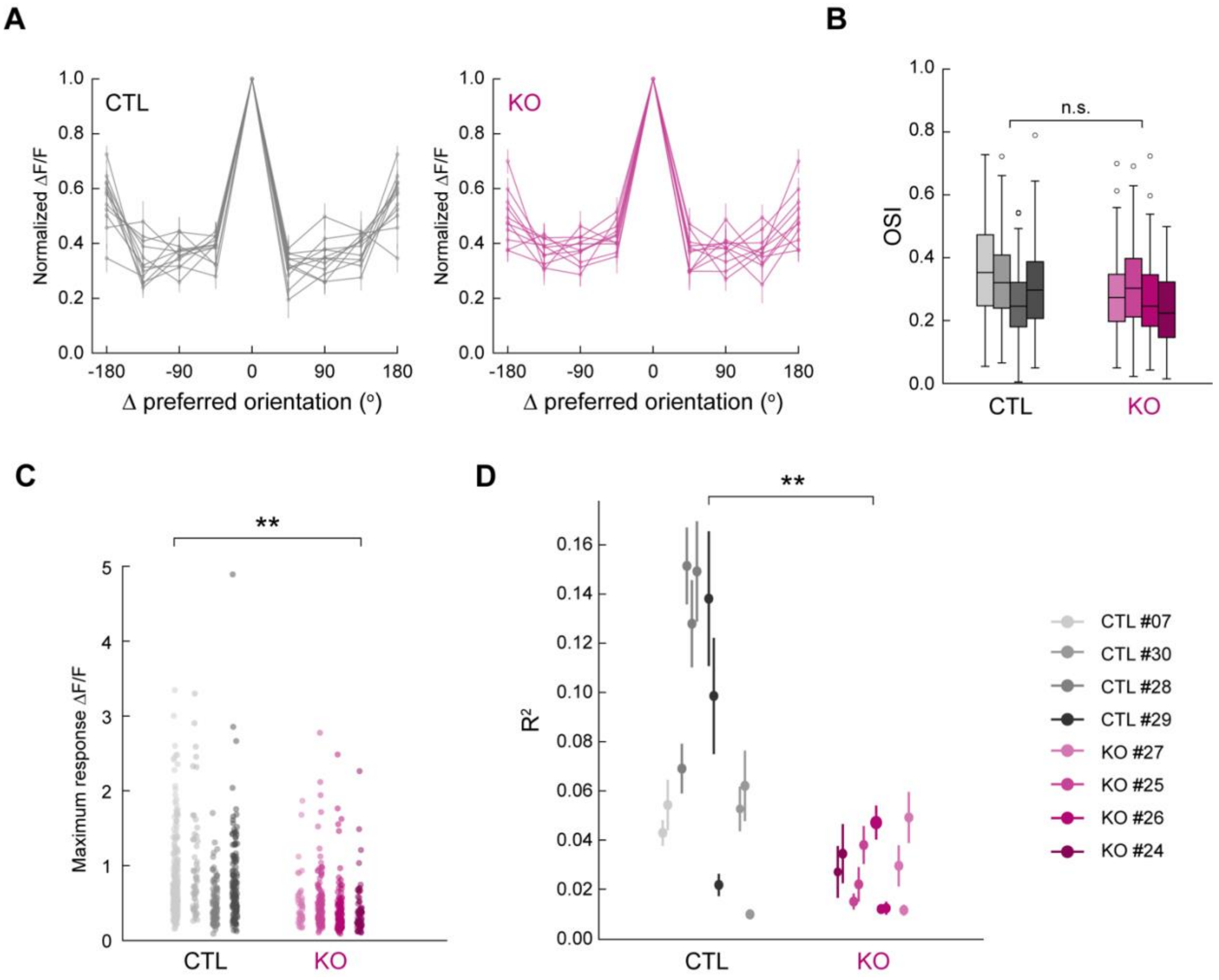


**Figure S5. Comparisons of visual responses of single neurons to drifting gratings** (A) Average tuning curves of neurons by session in each group ($n_{control}$ = 12 sessions, 4 mice, $n_{Gat3\ KO}$ = 11 sessions, 4 mice, error bars = SEM). (B) OSI distribution of neurons for each animal ($n_{control}$ = 4 mice, $n_{Gat3\ KO}$ = 4 mice, n.s., $p = 0.183$, LME t-stats, error bars = SEM). (C) Maximum response magnitudes of visually responsive neurons to their preferred gratings compared by animal ($n_{control}$ = 4 mice, $n_{Gat3\ KO}$ = 4 mice, **, $p < 0.01$, LME t-stats). (D) Average $R^2$ of neurons for single neuron encoding of drifting gratings and behavioral variables by sessions ($n_{control}$ = 12 sessions, 4 mice, $n_{Gat3\ KO}$ = 11 sessions, 4 mice, **, $p < 0.01$, LME t-stats, error bars = SEM).

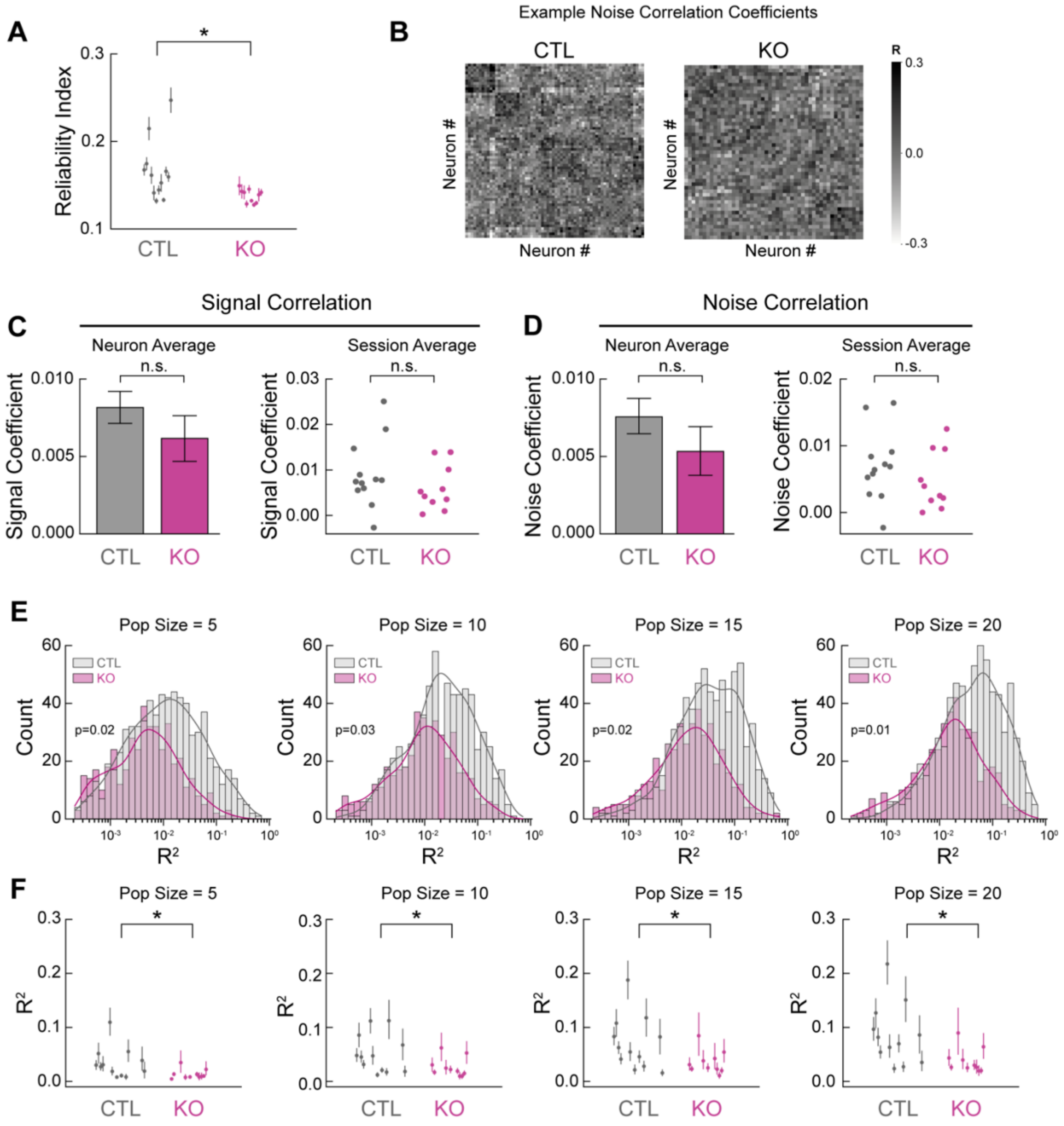


**Figure S6. Comparisons of neuronal responses to natural movies** (A) Average reliability indices of neurons per session ($n_{control}$ = 12 sessions, 4 mice, $n_{Gat3\ KO}$ = 10 sessions, 4 mice, *, $p < 0.05$, LME t-stats, error bars = SEM). (B) Representative noise correlation coefficient matrices. (C) Average signal correlation coefficients between pairs of neurons ($n_{control}$ = 24929 pairs, $n_{Gat3\ KO}$ = 10110 pairs, $n_{control}$ = 12 sessions, 4 mice, $n_{Gat3\ KO}$ = 10 sessions, 4 mice, n.s., $p = 0.741$, LME t-stats, error bars = SEM). (D) Average noise correlation coefficients between pairs of neurons ($n_{control}$ = 24929 pairs, $n_{Gat3\ KO}$ = 10110 pairs, $n_{control}$ = 12 sessions, 4 mice, $n_{Gat3\ KO}$ = 10 sessions, 4 mice, n.s., $p = 0.1349$, LME t-stats, error bars = SEM). (E) $R^2$ distribution of population activity encoding GLM model performance as a function of different population sizes of neurons used for model training. (F) Average $R^2$ values per session and per animal for each population size ($n_{control}$ = 12 sessions, 4 mice, $n_{Gat3\ KO}$ = 10 sessions, 4 mice, *, $p_{5\text{-}20} < 0.05$ each, LME t-stats, error bars = SEM).
